# Developing a novel recurrent neural network architecture with fewer parameters and good learning performance

**DOI:** 10.1101/2020.04.08.031484

**Authors:** Kazunori D Yamada, Fangzhou Lin, Tsukasa Nakamura

## Abstract

Recurrent neural networks (RNNs) are among the most promising of the many artificial intelligence techniques now under development, showing great potential for memory, interaction, and linguistic understanding. Among the more sophisticated RNNs are long short-term memory (LSTM) and gated recurrent units (GRUs), which emulate animal brain behavior; these methods yield superior memory and learning speed because of the excellent core structure of their architectures. In this study, we attempted to make further improvements in core structure and develop a novel, compact architecture with a high learning speed. We stochastically generated 30000 RNN architectures, evaluated their performance, and selected the one most capable of memorizing long contexts with relatively few parameters. This RNN, YamRNN, had fewer parameters than LSTM and GRU by a factor of two-thirds or better and reduced the time required to achieve the same learning performance on a sequence classification task as LSTM and GRU by 80% at maximum. This novel RNN architecture is expected to be useful for addressing problems such as predictions and analyses on contextual data and also suggests that there is room for the development of better architectures.

## 1 Introduction

Studies on neural networks and deep learning are currently expanding worldwide. The boom stems from the fact that neural networks have succeeded in realizing various complex tasks like processing images, understanding conversations, and writing sentences. The ability to understand conversations and write sentences is derived from the ability to understand contextual information. This is required for artificial general intelligence, which is an important goal of artificial intelligence research. Thus, methods to realize these abilities will be continuously developed and improved in the future. In the context of neural networks, these abilities are realized by recurrent neural networks (RNNs). The term RNN originally denoted a neural network that has recurrent loops in its structure. Various RNNs have been developed so far, including Elman networks (ENs) [1], echo state networks (ESNs), Boltzmann machines, long short-term memories (LSTMs) [2, 3], and gated recurrent units (GRUs) [4]. The most basic RNN is the EN, which has only one simple recurrent loop in its architecture. The ESN has a unique and unusual feature; the values of its weight parameters do not change from the initial random values. Instead, an ESN changes connections between neurons in the learning process. LSTM and GRU are popular architectures. Like EN, both LSTM and GRU exhibit recurrent loops in their architectures and can understand and handle contextual information of the input sequence data.

According to previous studies, the performances of LSTM and GRU, including the ability to retain contextual information for a long time and the learning speed of contextual information, are superior to that of EN. LSTM and GRU exhibit better performance by introducing gate structures to the architecture, mimicking the behavior of animal brains. Theoretically, the memory of LSTM can last forever because of the complex core algorithm based on gate structures, although this may not be necessarily true when we apply it to real world problems. The essential difference between EN and sophisticated methods like LSTM and GRU is the sophisticated core algorithm of the architecture, which undoubtedly improves the performance of RNNs. However, we believe that LSTM and GRU are not the best architectures for RNNs. At least, we cannot deny the possibility that there are better architectures than the LSTM and GRU because all possible architectures have not yet been examined; of course, the topological space of core algorithms is too large to examine all instances. In the present study, we developed a novel RNN architecture, showing better performance than some more sophisticated architectures. While developing a novel RNN algorithm, we focused on the compactness of architectures. Currently, many important advances of artificial intelligence are actually derived from deep learning, in which various neural network layers are stacked, thus increasing the number of parameters to achieve higher expressive power. This does indeed improve the performance of neural networks, and, in principle, we can increase the number of parameters indefinitely in this way. However, this is impossible in practice, as the physical limit to improving computer performance has nearly been reached [5]. Soon it will no longer be possible to increase the number of transistors on a microprocessor chip by reducing the size of transistors. From this viewpoint, we believe that a more compact architecture is better than a larger one if we can realize convergence performance with the compact architectures that is equivalent to that of the larger architectures, because a compact architecture has a lower computational cost [6]. In addition to such space constraints, the compact RNN structure also has an advantage in the computation time of the learning process. Generally, the inference process of an RNN with a large parameter size incurs a large computational cost. We expect that learning time could be reduced by using compact RNNs. In this study, based on the above ideas, we attempted to design a more compact architecture than GRU and LSTM. The objective of this study is to develop a compact RNN that has a shorter learning time than that of the existing large RNNs. The minimum requirements of the architecture developed in this study are as follows: compared to existing RNNs, the architecture has at least the same performance in terms of holding contextual information and has a high speed of learning based on a compact architecture.

In this study, we attempted to develop an architecture with good performance by generating candidate architectures randomly, evaluating them on a training dataset, and choosing the best one. Several studies based on such a concept can be found in the literature. For example, several studies mutated the core algorithm of LSTM or GRU and investigated the performance of the newly generated RNNs through various experiments [7, 8]. However, the number of core algorithms generated in those studies was limited. Because the new architectures were essentially variations on old ones, their performance was not always superior to that of existing RNNs; the search space was not sufficiently large. In addition, the studies did not set any restrictions on the new architectures; for example, they did not focus on the compactness of the core algorithm. On the other hand, several studies have indeed focused on this. For example, one study developed an RNN, simplified LSTM (S-LSTM), by reducing the number of gates from LSTM [9]. In other studies, minimal gated units (MGUs), embedded gated recurrent units (eGRUs), and simple gated units (SGUs) reduced a gate from a GRU [10–12]. In addition to the modifications of LSTM and GRU, the most basic RNN, EN, was also modified; independently recurrent neural network (IndRNN) changed a connection between the internal parameter variables of EN [13]. These compact RNNs were rationally designed based on modifying the existing RNNs. In the current study, by contrast, the topology of the core algorithm was randomly generated without any assumptions such as starting from the existing RNNs or modifying them. In our study, to generate compact architectures, the number of parameters in the core algorithms was restricted. The LSTM and GRU algorithms described in the next section (“Recurrent neural networks”) use four equations from **v_1_** to **v_4_** and three equations from **v_1_** to **v_3_** in their structures. In this study, we restricted the number of equations to two: **v_1_** and **v_2_**. Finally, the developed architecture, YamRNN (<u>Ya</u>mada <u>m</u>odified <u>RNN</u>), was evaluated on several test datasets and its properties, especially the performance of learning speed of contextual information, were investigated.

## 2 Recurrent neural networks

Among the RNNs that can incorporate contextual information of input data, the simplest architecture is EN [1]. EN is formulated by the following equation:

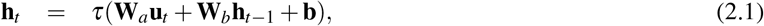

where **u**_*t*_, **h**_*t*_, +, ***τ***, **W**, and **b** denote an input vector for the RNN layer at timestep *t*, an output vector of timestep *t* (= internal memory vector of timestep *t*), an operator for the summation of matrices, a hyperbolic tangent function, a weight matrix, and a bias vector to be optimized. Generally, RNNs that can memorize contextual input sequences possess an internal memory vector **h**_*t*_ at each timestep. The vector is an output from an RNN unit, and at the same time, is maintained as a memory of the previous context. Although EN can memorize contextual data of input sequences by this mechanism, its performance is limited, mainly because of learning inefficiencies caused by the gradient vanishing problem. LSTM was developed to overcome this. LSTM has become one of the most famous RNN architectures [2]. The LSTM mimics the behavior of animal brains to improve performance in holding contextual information and in learning efficiency. The architecture of LSTM used in this study is defined by the following equations [3]:

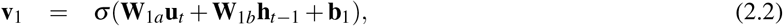

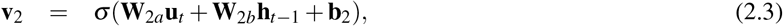

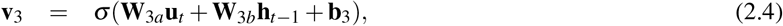

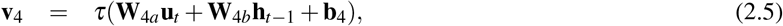

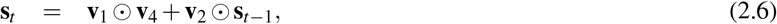

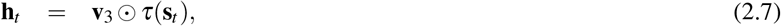

where **v**, **s**_*t*_, ⊙, and ***σ*** denote a temporal vector of the calculations, a unit for constant error at timestep *t*, an Hadamard product operator, and a sigmoid function. Equations 2.2 to 2.4 represent calculations of an input gate, a forget gate, and an output gate. Because the values of **v**_1_, **v_2_**, and **v_3_** range from 0 to 1, these vectors can also be considered gates when used as operands of the Hadamard product in equations 2.6 and 2.7. For example, if **v_1_** = **0**, then **v_1_** ⊙ **v_4_** = **0**, and the gate is closed; if **v_1_** = **1**, then **v_1_** ⊙ **v_4_** = **v_4_** and the gate is open. Equation 2.6 denotes the constant error carousel, which prevents the gradient from becoming too small or large. The final equation 2.7 denotes the memory vector, which is also used as an output vector from the LSTM cell. Using this memory vector, LSTM can memorize the previous input data and achieve the ability of reading and interpreting contextual data.

The other popular architecture, GRU, is newer than LSTM [4]. The architecture of GRU is defined by the following equations:

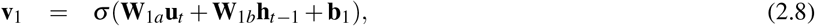

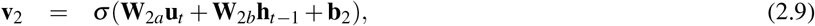

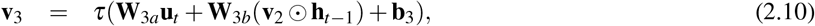

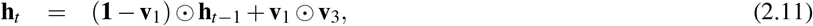

where **1** denotes the matrix (vector) of ones. There are a few differences between the core algorithms of LSTM and GRU. The first difference is the number of parameters used in the architectures. The core algorithm of LSTM has four intermediate vectors represented by **v** in equations 2.2 to 2.5. In the equations, there are a total of eight weight parameter matrices and four bias vectors. In contrast, GRU has three intermediate vectors, and the equations of the vectors have a total of six weight parameter matrices and three bias vectors. Thus, the number of parameters of GRU is 75% that of LSTM. The second difference between the architectures is the topology of the variables in the core algorithm. For example, equations 2.2 to 2.4 define the gate structure in LSTM, and they are utilized to calculate only the memory vectors (**s**_*t*_ and **h**_*t*_). By contrast, the gate structures are used to generate both an intermediate vector (**v_3_**) and a memory vector (**h**_*t*_) in GRU. In addition, the equations or connections between variables to generate the output vector (equations 2.7 and 2.11) in LSTM and GRU are largely different from each other. Collectively, these differences result in different performances of RNNs.

In addition to these popular RNNs, there are RNNs of compact architecture. The architecture of S-LSTM is defined by the following equations:

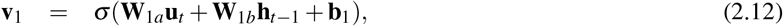

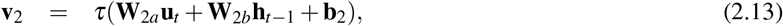

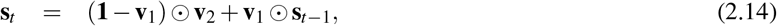

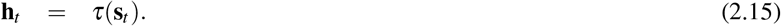

This architecture is quite similar to LSTM. It removes the output gates and replaces the input gate by the forget gate in the form of **1** – **v_1_**. On the other hand, the architecture of MGU is defined by the following equations:

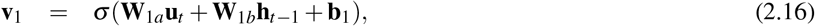

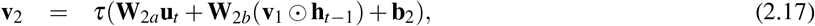

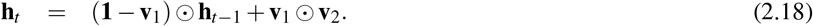

This architecture is quite similar to GRU. The architecture of eGRU is defined by the following equations:

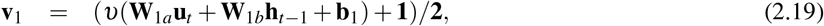

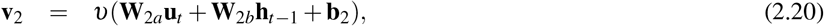

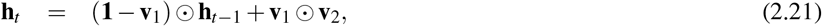

where *υ*, /, and **2** denote a softsign function, elementwise division, and the matrix (vector) of twos, respectively. The architecture of SGU is defined by the following equations:

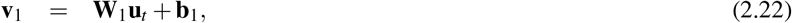

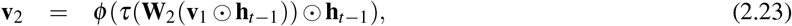

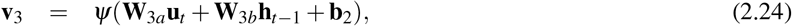

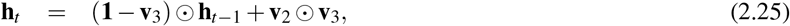

where *ϕ* and *ψ* denote a softplus and a hard sigmoid function, respectively. The parameters of these architectures include four weight parameters and two bias parameters. Thus, the number of parameters of the compact architectures is 50% that of LSTM. Further, the architecture of IndRNN is defined by the following equation:

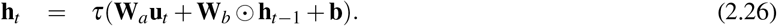

The architecture of IndRNN is quite similar to EN; these architectures do not contain any gate structures like the sophisticated RNNs. The parameters of the architecture include two weight parameters and a bias parameter. Thus, the number of parameters of IndRNN (and EN) is 25% that of LSTM.

## 3 Methods

### 3.1 Generation of RNN architectures

Herein, we generated various core algorithms of RNNs at random. In the generation phase, all stochastic processing followed a uniform distribution. The core algorithms were generated in a completely random way, except that they were constrained to meet the minimum requirements of having at least two weight parameters and a recurrent loop. The RNNs were generated according to the following rules:

1. The architecture has to include intermediate vectors, **v**_1_ and **v**_2_.
2. **v**_2_ has to be calculated after **v**_1_ is calculated.
3. The memory vector of the next timestep, **h**_*t*+**1**_, has to be calculated after **v_2_** is calculated.
4. The calculation of **v**_1_ and **v**_2_ must at least include a matrix product between the weight parameter matrix **W** and the input vector, **u**_*t*_ or the memory vector of the current timestep, **h**_*t*_. The bias parameter vector, **b**, is stochastically included.
5. To calculate **v**_2_, it is decided stochastically whether **v**_1_ is used. If **v**_1_ is used, **v**_1_ is transformed to **u**_*t*_ or **h**_*t*_ using any operator. The operators include summation (+), subtraction (−), and Hadamard product (⊙) of vectors. The operator is stochastically chosen.
6. The equations to calculate **v_1_** and **v_2_** are required to include a sigmoid (***σ***) or hyperbolic tangent (***τ***) as an activation function.
7. **h**_*t*+1_ is calculated using **v_1_**, **v_2_**, **h**_*t*_, and operators. However, at this time, if **h**_*t*_ has already been used to calculate **v_1_** or **v_2_**, it is not necessarily used again for the calculation of **h**_*t*+1_; whether or not to use it is decided stochastically. Similarly, if **v_1_** has already been used for the calculation of **v_2_**, the decision whether or not to use it again for the calculation of **h**_*t*+1_ is made stochastically.
8. As operators to calculate **h**_*t*+1_, summation (+), subtraction (−), Hadamard product (⊙) of matrices, and matrix product (×) are randomly chosen when variables **u**, **v_1_**, **v_2_**, or **h**_*t*_ are stochastically chosen. In addition, each variable can be subtracted from a vector of ones (**1**−) and can be squared in an elementwise manner. It is stochastically decided whether these operations are executed.
9. In the calculation of **h**_*t*+1_, the variables **u**, **v_1_**, **v_2_**, or **h**_*t*_ can be repeatedly selected without any restrictions. The recruitment of the variables is stochastically terminated.

Based on these rules, we stochastically generated 30000 architectures. We confirmed that each architecture had a unique topology and was different from the other architectures.

### 3.2 Dataset

#### 3.2.1 Screening and validation dataset

As a screening dataset for the generated RNNs, we utilized the simplest and most general dataset we could in order to reduce the possibility of biased results due to specific datasets. Thus, we generated a dataset including 10 instances. Each instance had a length of 402 timesteps and consisted of a sequence of integers between 0 and 399, one integer per timestep. The integer corresponding to timestep 0 of instance *n* (where *n* = 0.1, …, 9) was chosen to be simply *n*. The same value *n* was also assigned to timestep 401 of instance *n*, so the first element of each instance was the same as the last. The integers corresponding to timesteps 1 to 400 did not depend on the instance but obeyed the simple rule: timestep *t* corresponds to the integer *t* – 1, where *t* = 1, 2, …, 400. The difference between each instance in the dataset was therefore only the initial and final elements. Thus, if we face a problem in which the predictor of RNNs predicts a value of timestep *t* + 1 by accepting a value of timestep *t* as an input vector, and if the predictor succeeds in predicting the final value, the predictor can at least hold 400 contextual elements. Therefore, this dataset is the most general benchmark dataset for the evaluation of RNNs that can memorize contextual information.

As a validation dataset for screened RNNs, we utilized the Modified National Institute of Standards and Technology (MNIST) [14] dataset. The MNIST is a famous benchmark dataset for classification of handwritten digits. Because instances in the dataset were image data with 28 rows and 28 columns, we regarded an instance as a sequential datum with a length of 28, in which each element consisted of a 28-dimensional vector. The number of instances in the dataset is 70000, and we utilized all of them for the validation process.

#### 3.2.2 Test dataset

As the test dataset for the developed RNN and other existing RNNs, we utilized S&P 500 stock data obtained from Kaggle (https://www.kaggle.com/camnugent/sandp500), which contained historical stock data for all S&P 500 companies for five years. From the original dataset, we removed nine instances including “BHF”, “DHR”, “ES”, “FTV”, “O”, “REGN”, “UA”, “VRTX” and “WRX” because they contained missing values. The instances in the dataset are time series data. Each timestep in an instance corresponds to a five-dimensional vector in which each element can take the values “open”, “high”, “low”, “close”, and “volume”. From the original dataset, in which an instance of each company has a sequential vector structure with a length of 1259, we randomly extracted data of length 100, 300, and 500 for the test dataset by not allowing any overlap between sequences. We named these datasets test dataset 1 (SP500–100), test dataset 2 (SP500–300), and test dataset 3 (SP500–500), respectively; each dataset included 1340, 851, and 631 instances, respectively. The values of each instance were standardized to z-values. For the dataset, RNNs predicted “up” or “down” of “close” values by reading the previous contextual information before the date of prediction. We chose this dataset because predicting stock price is known to be difficult, and thus we could easily recognize differences of learning performance between the architectures. Moreover, we could change the sequence length of instances freely (100, 300, and 500) to measure and compare the performance of different RNNs as a function of sequence length. In addition to the datasets, we also utilized the IMDb movie review sentiment classification dataset, which is one of the most famous benchmark datasets used for evaluating the performance of RNNs [15]. The dataset includes 25000 comments of movie-reviews written in English. The length of each comment differs, with the lengths of the shortest and longest comments being 11 and 2494, respectively. The dataset comprises a total of 88586 unique words. All comments are linked to sentiments such as “positive” or “negative” feeling. For the dataset, RNNs were engaged in the sentiment analysis and predict “positive” or “negative” for the comment values by reading the contextual information of each comment. In the study, we named the dataset test dataset 4 (IMDb). Further, we utilized the Reuters newswire classification dataset, provided by TensorFlow [16] as another test dataset. Similar to the IMDb, this dataset includes sentences written in English; there are 8982 instances, with lengths ranging from 13 to 2376. This dataset comprises a total of 30981 unique words, and the sentences are linked to 46 topics. In this study, we named the dataset test dataset 5 (Reuters). For the dataset, RNNs were engaged in a document classification task and predict the topic to which the sentences belong by reading the contextual information of each sentence. We chose the two datasets in order to evaluate the performance of RNNs on datasets comprising variable-length data, because one of the merits of using RNNs is that they can competently process variable-length data, and evaluating RNNs on the datasets will help us to understand the usefulness of RNNs under a situation that is similar to the real world.

### 3.3 Screening and validation of architectures

The 30000 newly generated architectures were screened using the screening dataset. The size of the weight parameter was fixed as 24576, i.e., the product of the size of the bias parameter (512) and the length of the input vector (48), as explained below. In addition to the RNN layer, only one fully connected layer was added to produce an output vector of a suitable size for the given classification problem. The input value, which was an integer from 0 to 399, was encoded to a floating value vector of 48-dimensions. All weight and bias parameters were randomly initialized by the uniform distribution *U*(−0.2, 0.2). The size of the minibatch was fixed as the instance number, 10, and parameter update was conducted by the batch update method. As an optimizer of the gradient descent method, Adam was utilized [17]. In the process, for every candidate architecture, the value of the prediction accuracy was recorded for every epoch. If the accuracy was serially recorded as 1.0 for 10 epochs, the architecture was regarded as having “succeeded” and the epoch was recorded. This computation was repeated five times for each architecture. If the epoch exceeded 50000 times, the architecture was regarded as having “not succeeded” and the computation was stopped.

For the validation process, almost all settings were the same as those in the screening process. The differences were the encoding process of the input vector and the size of the minibatch. In the validation process, because the value of the input vector in the MNIST dataset was not an integer (categorical) value, the encoding process was not performed. The size of the minibatch was set to 1000. In addition, the threshold of the prediction accuracy was set to 0.999 instead of 1.0, and the computation was repeated 10 times because the computational burden was not so heavy for the problem.

### 3.4 Investigation and test conditions

The performance of YamRNN, the architecture we developed, was investigated from various aspects. It was examined on the screening dataset by changing the hyperparameter settings such as the size of weight and bias parameters, the minimum and maximum value of uniform distribution for parameter initialization, and the number of instances in the dataset. More details are provided in the next section (“Results and discussion”) for each investigation.

The performance of YamRNN was compared with that of existing RNNs on the test datasets. In the experiments on test datasets 1–3, the size of the weight parameter was changed from 256 × 5 to 1024 × 5 with an increment of 256 × 5, where “5” comes from the five values “open”, “high”, “low”, “close”, and “volume” of an element of an instance in the test dataset. The size of the bias parameter was changed from 256 to 1024 with an increment of 256. In addition to the RNN layer, only one fully connected layer was added to produce the output vector of a suitable size for the classification problem, i.e., two (“up” or “down”). On the other hand, in the experiments on test datasets 4 (IMDb) and 5 (Reuters), the size of the weight parameter was varied using values of 32 × 8, 64 × 8, 128 × 8, and 256 × 8, where “8” is the length of the word-embedded vector, because we embedded each word as an eight-dimension vector. Correspondingly, the size of the bias parameter was varied using values of 32, 64, 128, and 256, respectively. The sizes of the parameters were smaller than those in the first three tests owing to the limitation of our computational environment. In addition to the RNN layer, only one fully connected layer was added to produce an output vector of a suitable size for the classification problem, i.e., two (“positive” or “negative”) for IMDb or 46 (topics) for Reuters. All weight and bias parameters were randomly initialized by the uniform distribution *U*(−0.2, 0.2). The size of the minibatch was set to 300. As an optimizer of the gradient descent method, Adam was utilized. In the process, the value of the prediction accuracy was recorded for every epoch. If the accuracy was serially recorded over 10 epochs as 0.7 for test datasets 1-3 and 5, and 0.9 for test dataset 4, the trial was regarded as having “succeeded”, and the epoch was recorded. For the first three experiments and the fifth experiment, we set the threshold value to 0.7 because of the problem’s high level of difficulty; it was difficult for any architecture to reach higher accuracy than approximately 0.7. In contrast, for the experiments on IMDb, we set the threshold value to 0.9 because the difficulty of the problems was low; this value is reasonable as we confirmed the simplicity of the dataset in various published studies that utilized the dataset [10, 12, 18]. The computation was repeated 10 times for each test. If the epochs exceeded 50000 times on SP500–100, SP500–300, and SP500–500 or 500 times on IMDb and Reuters, the trial was regarded as having “not succeeded”, and the computation was stopped. The number of successful trials was counted, and, in addition to the final epoch, the actual computation time (seconds) was recorded.

### 3.5 Computation environment

As a framework to implement the neural networks, we used Theano version 1.0.4 [19] with CUDA version 10.0 (NVIDIA) and cuDNN version 7.5 (NVIDIA), and TensorFlow version 2.1.0 with CUDA version 10.1 and cuDNN version 7.6 for the sentiment analysis on IMDb and document classification on Reuters. All calculations were performed on the NIG supercomputer at ROIS National Institute of Genetics with Tesla V100 (NVIDIA) as a GPU accelerator.

## 4 Results and discussion

### 4.1 Development of a novel architecture

In this study, we screened 30000 newly generated RNN architectures by benchmarking them on the screening dataset. Before the screening process, we evaluated the performances of existing method, such as LSTM and GRU, on the dataset. The results are shown in Figure 2. The vertical axis includes the number of epochs for each RNN predictor to reach the threshold accuracy (=1.0). Thus, a predictor with a small value on the vertical axis denoted that the predictor had a high learning speed for the input sequence. As seen from the data shown in Figure 2 the learning speed of GRU was found to be superior, which is consistent with the literature data [10]. Longer contextual information requires more time to learn. From these results, we could elucidate the approximate time required for learning with the RNNs. We tried to screen the architectures on as long a sequence dataset as possible; by considering the expected computational burden and the performance of our computer environment, we set the length of the context used for the screening process to 400. In addition, we set the maximum limit of screening time to 50000 epochs to save computational resources. The objective of the study was to develop an RNN that had a compact core algorithm (smaller number of parameters) and thus showed good learning speed with descent memorizing performance. First, we generated 30000 RNN architectures, based on the procedure described in the “Methods” section. The algorithms of each architecture were unique. Next, the performance of the architectures was examined on the screening dataset. Thus, 21148 out of 30000 architectures did not work as normal RNNs because they did not return valid values of cost and prediction accuracy. Only 579 of the remaining 8852 RNNs passed all the other screening criteria.

**Fig. 1.**
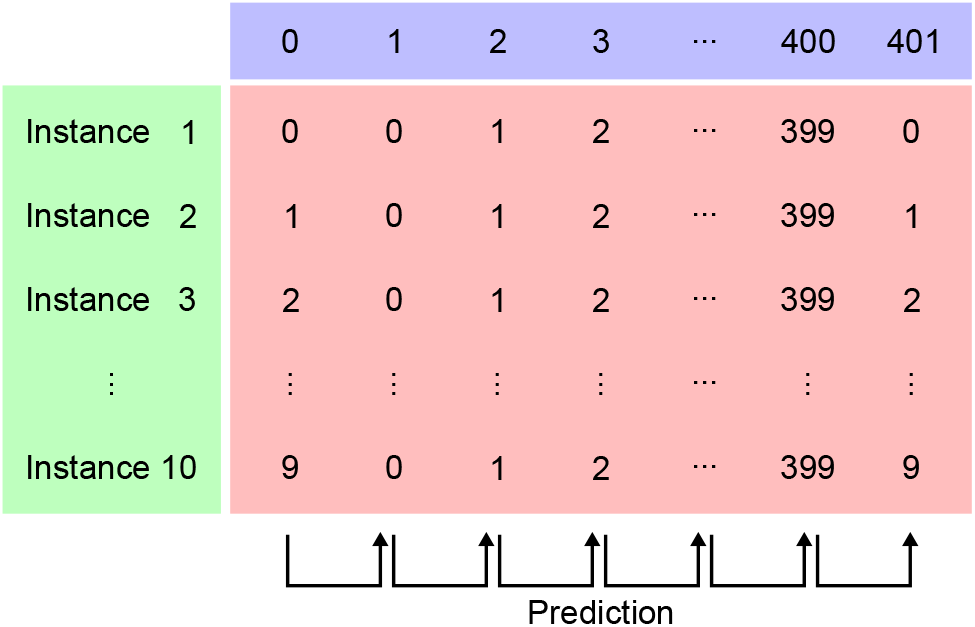
Illustration of the screening dataset. The numbers in the purple box denote the timestep *t*. In each instance, the initial and final elements are the same. The middle elements are common for all instances. The value of the next timestep is predicted using RNNs based on the value of the previous output as the input.

**Fig. 2.**
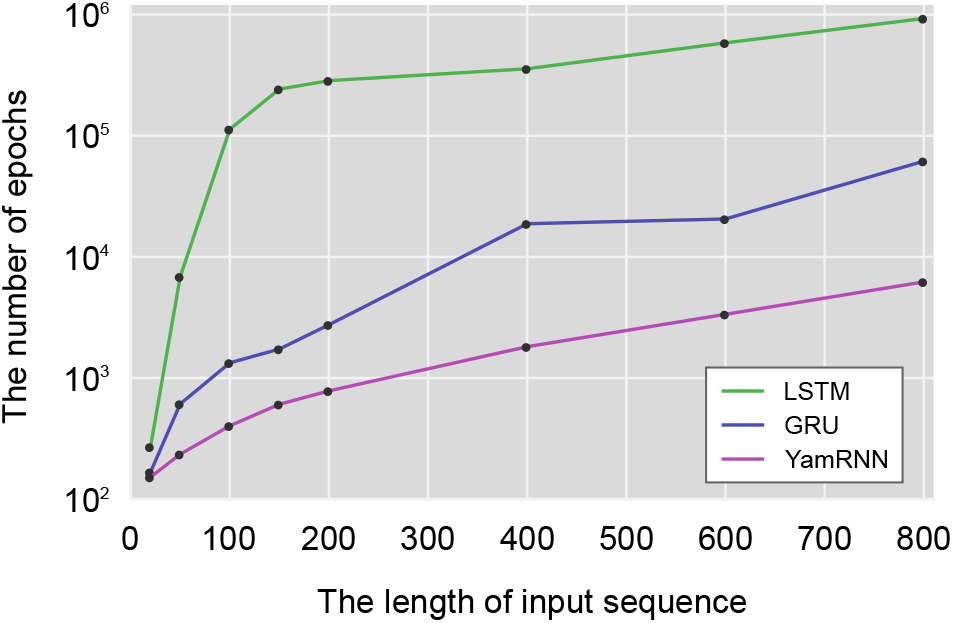
Comparison of learning speeds of RNNs on the screening dataset. The horizontal axis represents the size of units of weight and bias parameters for each RNN, and the vertical axis represents the number of epochs for the RNN predictors to serially reach the threshold accuracy (= 1.0) 10 times.

Next, we proceeded to a second screening process, which we call the validation process. As described in the “Methods” section, we utilized the MNIST dataset as contextual data consisting of 28 timesteps. In contrast to the screening dataset, which was a simple simulation dataset of longer contexts, the MNIST dataset had shorter contextual information, was derived from real world data, and included numerous instances. By using different validation datasets, we expected the RNNs remaining after two screenings to show more universal performance. As in the first screening, we first examined the performance of LSTM and GRU on the dataset before conducting the validation experiment. As shown in Table 1, the mean numbers of epochs for LSTM and GRU to reach the threshold accuracy (= 0.999) were 30.1 and 25.3, respectively. Again as in the previous experiment on the screening dataset, the learning speed of GRU was superior to that of LSTM. Subsequently, we conducted a validation experiment. Although none of the RNNs that went through the validation process output an invalid cost value or accuracy, only 304 architectures completed the computation in 50000 epochs. Because the number of epochs of GRU on the dataset was approximately 25, first we attempted to find architectures whose number of epochs was less than 25. At this time, we sorted the candidate RNNs in ascending order of the cost value. However, none of the surviving architectures recorded a number of epochs less than that of GRU. Therefore, we changed the threshold to 30 and further to 40; finally, we found an RNN architecture that recorded 36.0 as the number of epochs (Table 1). We assigned the name “YamRNN” to the one RNN architecture that succeeded in our study. The core algorithms of YamRNN were defined using the following equations:

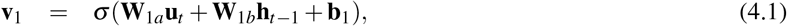

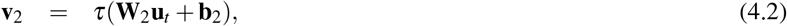

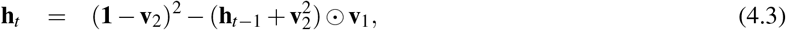

where **x**^2^ denotes **x** ⊙ **x**. The number of parameters of YamRNN is compared with the existing architectures in Table 2. As described, YamRNN has three weight and two bias parameters. The number of parameters of YamRNN is less than 50% that of LSTM and 67% that of GRU; it is also smaller than that of existing compact architectures such as S-LSTM, MGU, eGRU, and SGU, but not smaller than that of the primitive RNNs without gate structures such as EN and IndRNN. Equation 4.1 includes a sigmoid function, which can be considered to work as a gate, like those of existing RNNs. Next, we benchmarked YamRNN on the screening dataset by changing the length of the input sequences. The performance of YamRNN on the dataset is shown by a purple line in Figure 2. As shown, the learning speed of YamRNN was superior to that of LSTM and GRU for any input length. This tendency was especially prominent for longer input sequences, and the learning speed of YamRNN was approximately 150 and 9.92 times that of LSTM and GRU, respectively, on the dataset in which the length of the input sequence was 800.

**Table 1.**
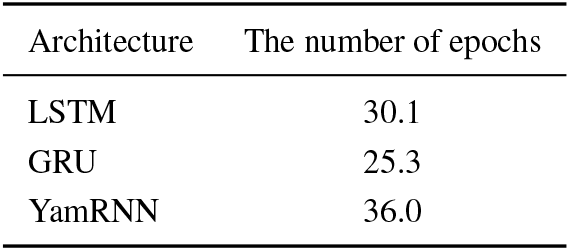
Number of epochs for RNNs to reach the threshold accuracy on the validation dataset. The value was the mean of 10 trials. Three significant digits are displayed.

| Architecture | The number of epochs |
| --- | --- |
| LSTM | 30.1 |
| GRU | 25.3 |
| YamRNN | 36.0 |

**Table 2.**
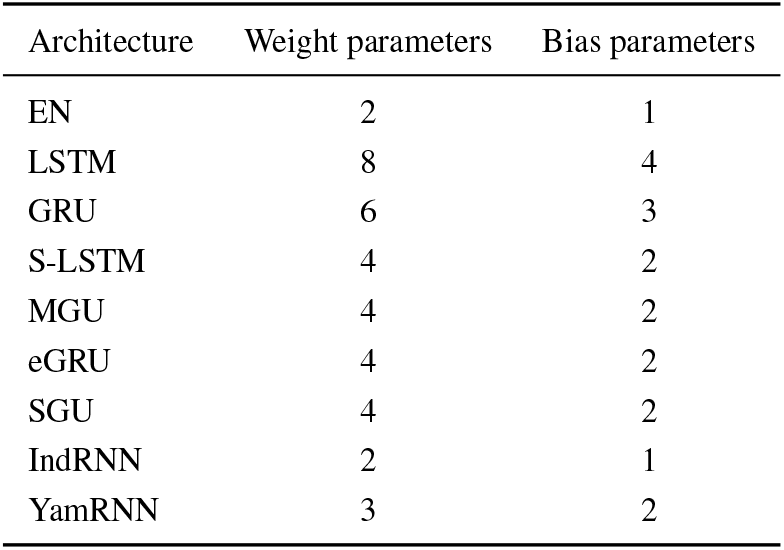
Comparison of the number of weight and bias parameters in the equations of each RNN

| Architecture | Weight parameters | Bias parameters |
| --- | --- | --- |
| EN | 2 | 1 |
| LSTM | 8 | 4 |
| GRU | 6 | 3 |
| S-LSTM | 4 | 2 |
| MGU | 4 | 2 |
| eGRU | 4 | 2 |
| SGU | 4 | 2 |
| IndRNN | 2 | 1 |
| YamRNN | 3 | 2 |

### 4.2 Investigating the performance of YamRNN

Next, we investigated the performance of YamRNN by changing the hyperparameter settings, including the number of units of RNN layer, the parameter of uniform distribution, and the number of instances in a dataset. In the experiments, we benchmarked the performance of YamRNN on the datasets, which were similar to the screening dataset but had different input-sequence lengths: 50, 100, 200, 400, or 800. If the accuracy was recorded as 1.0, 10 times serially, computation was stopped and the computation time (seconds) was recorded. The conditions other than the changed hyperparameter settings in each experiment were the same as those of the first screening. First, we benchmarked the performance of YamRNN by changing the number of units in the layer of the architecture. The benchmark results are shown in Figure 3a, where the five columns of different colors denote the computation time of learning by YamRNN on datasets of sequence lengths 50, 100, 200, 400 and 800 respectively. As the length of the instances increased, larger computation times were required. In contrast, the learning speed increased as the number of units in the layer increased. Generally, a model with more parameters has more expressive power. In this experiment, the learning process was completed quickly because of the high expressive power, so the calculation time of the model with larger number of parameters could be reduced. This tendency was observed in the subsequent test experiments described in the next section. Next, we conducted benchmarking by changing the parameter of uniform distribution used to initialize the weight and bias parameter of models. As shown in Figure 3b, the computation time increased depending on the length of the instances in each dataset. Regarding the change of parameters of uniform distribution, when we set the distribution as *U*(−0.4, 0.4), the computation time on almost all datasets was minimized. This means the learning speed was maximized on the problem when the parameter of uniform distribution was set to 0.4. However, we must note that the parameter of uniform distribution is only a hyperparameter, and the result is not a universal setting; the value of 0.4 was just used for reference. Finally, we performed benchmarking by changing the size of the dataset. As shown in Figure 3c, when the number of instances in the dataset increased, the computation time also increased. This result is unsurprising, because the computation load naturally increases with increase of the dataset size, and the problem becomes more difficult depending on the dimension of the output vector, which is the same as the number of instances. The dimensions of the output vector on a dataset with 10 instances and 400 instances were 10 and 400, respectively. Collectively, by investigating the performance of YamRNN, we confirmed that the performance of the architecture changed proportionally depending on the change of parameters. Thus, the behavior of the architecture against parameter changes was considered to be normal.

**Fig. 3.**
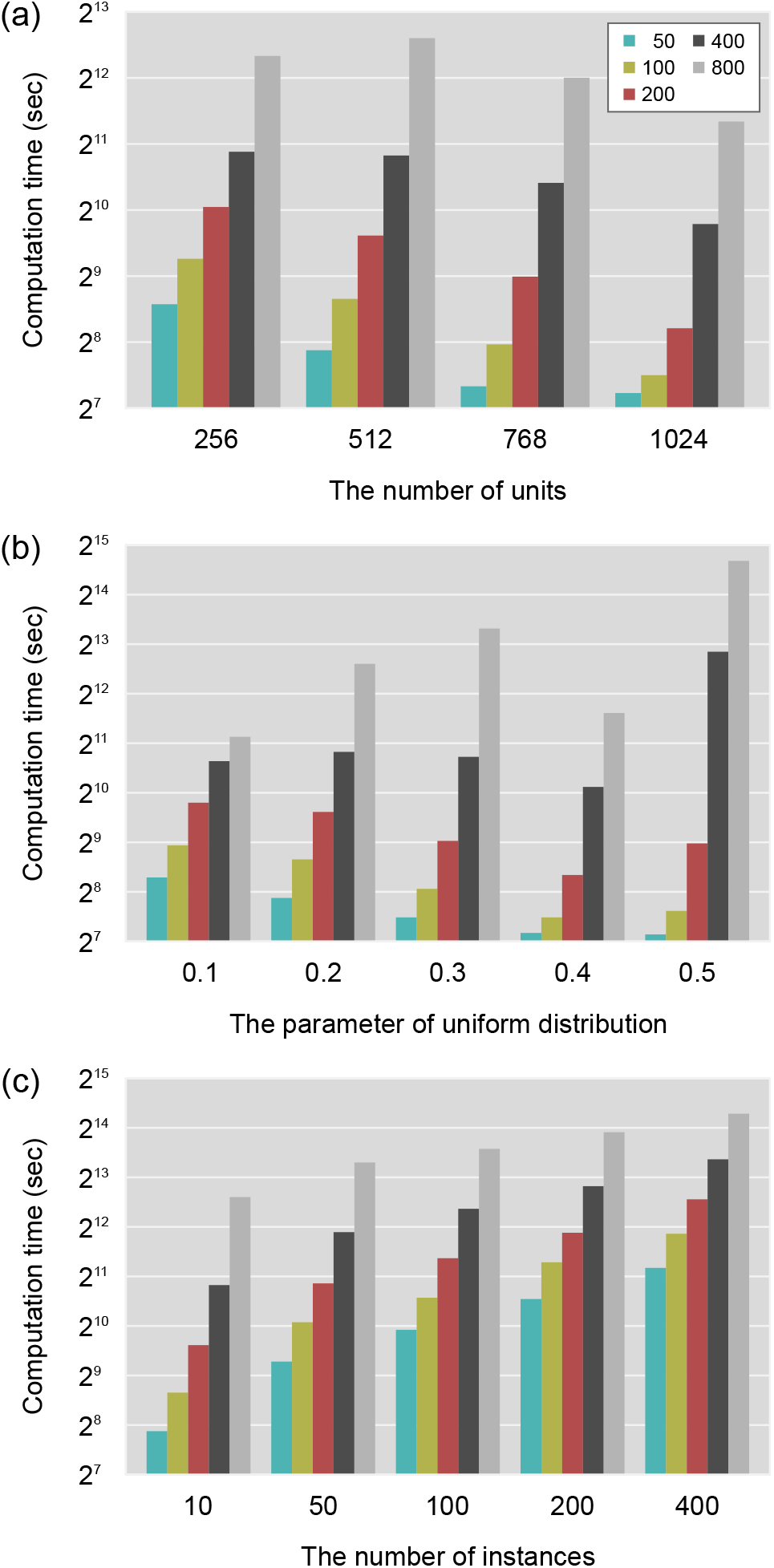
Comparison of learning speed of YamRNN under various conditions. The vertical axis represents the computation time for a model to reach threshold accuracy (= 1.0) 10 times serially on datasets, where instances in each dataset have different lengths from 50 to 800, represented by columns of different color. (a) The horizontal axis represents the number of units on weight and bias parameters. (b) The horizontal axis stands for the parameter of uniform distribution, by which the parameters of a model are initialized. (c) The horizontal axis represents the number of instances in the dataset.

### 4.3 Comparison to the existing methods on learning speed

Next, we compared the performance of YamRNN on the test datasets to that of existing architectures such as EN, LSTM, GRU, S-LSTM, MGU, eGRU, SGU, and IndRNN. As in the previous experiments, the criterion to evaluate the goodness of architectures was the learning speed, as reflected in the number of epochs and the actual computation time (seconds) to reach the threshold accuracy. As described in the “Methods” section, in the test, we computed the learning speed on the nine architectures by changing the number of units of the RNN from 256 to 1024 at an increment of 256 for test datasets 1–3, and 32, 64, 128, and 256 for test datasets 4 and 5. The number of times the computation was completed out of 10 trials was recorded. We judged the computation as completed when the prediction accuracy recorded the threshold accuracy (0.7 or 0.9) serially 10 times within the threshold number of epochs (50000 or 500). First, we conducted benchmarking on SP500–100, where the length of input sequences was 100. As shown in Table 3, many architectures except EN and IndRNN completed the computation within the threshold number of epochs on each trial. When the number of units was 256, GRU and MGU showed the best learning speed. If the number of parameters of LSTM, GRU and MGU was 512, and that of S-LSTM and eGRU was 768, the learning speed reached the best value; if the size was increased further, the performance deteriorated because the optimization phase of the increased parameters required more time. By contrast, the learning speed of YamRNN worsened with smaller sizes of parameters (256) but was excellent for larger sizes (768 and 1024). We could interpret the results as follows: YamRNN only achieved adequate expressive power at a certain parameter size. Actually, the number of parameters (both weight and bias) of YamRNN on 256 units was 4352, only about half that of GRU (8448). Collectively, the learning time required to achieve the same learning performance as that of LSTM and GRU, currently the most popular RNN, was shortened at maximum by 85% and 80% on the test dataset when the number of units was set to 1024. Meanwhile, the primitive RNNs, EN and IndRNN did not record a decent performance on any experiments, which is consistent with their performance reported in many studies on RNNs [2]. In addition to difference in the number of parameters, one of the remarkable differences between the primitive RNNs and the other RNNs examined in this study is the existence of the gate structures; the primitive RNNs do not have such structures. Although the number of parameters of YamRNN was less than that of the sophisticated RNNs, the performance of YamRNN, which has a gate structure, was good. From this perspective, the existence of the gate structures is an important factor impacting the performance of RNNs. Next, we conducted the same benchmarking on SP500–300, where the length of the input sequences was 300. As shown in Table 4, the problem was more difficult than the previous test because the success count was smaller and the number of epochs and computation time were larger than the previous results. As in the previous case, the performance of IndRNN was quite bad and at this time that of EN was worse than the previous results. This reflects the fact that EN could not process longer sequences owing to the learning inefficiencies caused by the gradient vanishing problem [2]. Consequently, LSTM and its derivative, S-LSTM, failed to complete the computation in most cases. At the same time, SGU, which is not a derivative of LSTM, also failed to complete the computation in many cases. Considering these facts, it is considered that the performance deterioration of these RNNs is not due to some property of LSTM but other properties of RNNs. In addition, GRU failed to complete the computation when the parameter size was 1024, possibly because the threshold time for the problem was not long enough for GRU to optimize parameters. As before, the learning speed of YamRNN was inferior to that of LSTM, GRU and MGU when the number of parameters was small (256). By contrast, if the parameter size was large, the learning speed of YamRNN was superior to those of the other methods. Surprisingly, the learning speed of YamRNN did not diminish despite the increase in the number of parameters. This property was not seen in any other architecture and suggests YamRNN’s excellent potential. Next, we conducted benchmarking on SP500–500, where the length of the input sequences was 500. As shown in Table 5, the problem was more difficult than either of the other two tests. In addition to EN and IndRNN, neither LSTM nor S-LSTM succeeded in completing the computation on the tested parameter settings and even GRU and MGU, which had good performance on the previous two tests, could not finish the computation when the size of the parameter was large (768 and 1024). By contrast, YamRNN could complete the computation when the size of the parameter was large, although it failed to do so when the parameter size was small. This was once again because of the limited expressive power of YamRNN in the small parameter setting. When the parameter size was large, the learning speed of YamRNN was stably high. This tendency was one of the most outstanding features of YamRNN. Next, we conducted benchmarking on IMDb, with differing input sequence lengths. The most notable difference between this dataset and the previous test datasets is that this dataset is easier for learning; more specifically, features inside the dataset are elucidated more easily by RNNs than the previous test datasets. In fact, as shown in Table 6, IndRNN, which could not complete any trials in the previous tests, completed all the trials in this experiment. The performance of IndRNN improved with increasing parameter size, which is the same tendency that YamRNN uniquely showed in the previous tests. Similarly, the performance of MGU was also excellent in this experiment. However, the performance deteriorated with increasing parameter size, which is the same tendency as in the previous tests. Meanwhile, LSTM and MGU showed the highest learning speed when the parameter size was 64 and GRU showed the highest learning speed when the parameter size was 128, which is the median parameter size examined in the tests. Thus, the highest performance was not always achieved by RNNs with the largest or smallest parameters. Taking this observation into consideration, the optimal parameter size of each RNN would depend on the problem. We surmise that this is logical, which is why we search for the best hyperparameter when working with machine learning applications. As in the previous tests, the performance of YamRNN continued to improve with increasing parameter size; the performance of YamRNN with large parameter size was the second best among the RNNs following IndRNN. Although we could not examine the performance of YamRNN with larger parameters owing to the limitation of the current computation power, we estimate that the performance of YamRNN would continue to improve until it reached the optimal parameter size and the best performance if the parameter size could be further increased. This might also be true of SGU and IndRNN, but considering the previous test results, these RNNs would not always give stable performance. Next, we conducted benchmarking on Reuters with differing input sequence lengths. In contrast to the test on IMDb, which was a binary classification problem, the test on Reuters was a multi-class classification problem with 46 classes. As shown in Table 7, the performance of many RNNs, such as GRU, MGU, SGU, IndRNN, and YamRNN, continued to improve with increasing parameter size. As mentioned above, an optimal parameter size exists for each RNN for each problem and the optimal parameter of these RNNs for this problem would be larger than the sizes that we examined in the study. Although these RNNs had the same tendency with increasing parameter size, their performances differed completely; in this problem, the performance of YamRNN was good and was always the best among the RNNs. Although MGU had a good performance on several parameter settings in the previous problems, the performance of YamRNN was overwhelmingly better than that of MGU on this problem. Considering the computation time of each RNN, the difficulty of convergence on Reuters was clearly higher than that on IMDb. Similarly, the difficulty of convergence on SP500–100, SP500–300, and SP500–500 increased in this order. The performance of YamRNN was better on Reuters than IMDb; similarly, it was better on SP500–500 than SP500–100 or SP500–300. Taking this fact into consideration, the performance of YamRNN might be superior to other RNNs in more difficult and complex problems, although we would need more experiments on various datasets in order to prove this assertion. As shown, the learning speed of YamRNN was good and it at least converged within practical time when the parameter size was small. Furthermore, the learning speed was excellent when the parameter size was large, even though we examined it on more general benchmark datasets.

**Table 3.** Results on test dataset 1 (SP500–100), where the length of input sequences was 100. The number of epochs and computation time (seconds) is the mean value of the successful trials. The best values for each category are shown in bold. Three significant digits are displayed.

| The unit size of RNN layer | Success count out of 10 trials |  |  |  | Number of epochs |  |  |  | Computation time |  |  |  |
| --- | --- | --- | --- | --- | --- | --- | --- | --- | --- | --- | --- | --- |
|  | 256 | 512 | 768 | 1024 | 256 | 512 | 768 | 1024 | 256 | 512 | 768 | 1024 |
| EN | 4 | 7 | 4 | 0 | 7460 | 12900 | 15000 | – | 893 | 2100 | 2800 | – |
| LSTM | **10** | **10** | **10** | **10** | 2900 | 1430 | 2970 | 4520 | 1250 | 925 | 2120 | 3490 |
| GRU | **10** | **10** | **10** | **10** | 2160 | 1950 | 2140 | 4090 | 729 | 983 | 1220 | 2530 |
| S-LSTM | 9 | 9 | **10** | **10** | 4350 | 4940 | 3480 | 5750 | 1070 | 1040 | 1400 | 2500 |
| MGU | **10** | **10** | **10** | **10** | **1940** | **1170** | 4230 | 7830 | **491** | **421** | 1750 | 3480 |
| eGRU | **10** | **10** | **10** | **10** | 4570 | 3070 | 2910 | 5840 | 1080 | 1040 | 1130 | 2480 |
| SGU | 5 | **10** | **10** | **10** | 27200 | 13200 | 10300 | 9440 | 7410 | 5100 | 4540 | 4440 |
| IndRNN | 0 | 0 | 0 | 0 | – | – | – | – | – | – | – | – |
| YamRNN | **10** | **10** | **10** | **10** | 4240 | 1850 | **1690** | **1680** | 795 | 461 | **473** | **507** |

**Table 4.** Results on test dataset 2 (SP500–300), where the length of the input sequences is 300. The number of epochs and computation time (seconds) is the mean value of the successful trials. The best values on each category are shown in bold. Three significant digits are displayed.

| The unit size of RNN layer | Success count out of 10 trials |  |  |  | Number of epochs |  |  |  | Computation time |  |  |  |
| --- | --- | --- | --- | --- | --- | --- | --- | --- | --- | --- | --- | --- |
|  | 256 | 512 | 768 | 1024 | 256 | 512 | 768 | 1024 | 256 | 512 | 768 | 1024 |
| EN | 0 | 0 | 0 | 0 | – | – | – | – | – | – | – | – |
| LSTM | 4 | 2 | 3 | 0 | 11500 | 26500 | 42200 | – | 7180 | 25200 | 44400 | – |
| GRU | **10** | 9 | **10** | 3 | 7800 | 12800 | 31000 | 44700 | 3910 | 9520 | 26000 | 41000 |
| S-LSTM | 2 | 5 | 7 | 5 | 15900 | 24500 | 31900 | 44000 | 5750 | 12800 | 18900 | 28600 |
| MGU | **10** | 7 | 2 | 0 | **7780** | 25800 | 45400 | – | **2890** | 13800 | 27600 | – |
| eGRU | 8 | 9 | 7 | 7 | 17900 | 15800 | 23300 | 25900 | 6200 | 7910 | 13300 | 16400 |
| SGU | 0 | 0 | 5 | 9 | – | – | 42800 | 40100 | – | – | 28200 | 26800 |
| IndRNN | 0 | 0 | 0 | 0 | – | – | – | – | – | – | – | – |
| YamRNN | 7 | **10** | **10** | **10** | 19000 | **5730** | **4930** | **3950** | 5270 | **2120** | **2050** | **1770** |

**Table 5.** Results on test dataset 3 (SP500–500), where the length of the input sequences is 500. The number of epochs and computation time (seconds) are the mean values of the successful trials. The best values for each category are shown in bold. Three significant digits are displayed.

| The unit size of RNN layer | Success count out of 10 trials |  |  |  | Number of epochs |  |  |  | Computation time |  |  |  |
| --- | --- | --- | --- | --- | --- | --- | --- | --- | --- | --- | --- | --- |
|  | 256 | 512 | 768 | 1024 | 256 | 512 | 768 | 1024 | 256 | 512 | 768 | 1024 |
| EN | 0 | 0 | 0 | 0 | – | – | – | – | – | – | – | – |
| LSTM | 0 | 0 | 0 | 0 | – | – | – | – | – | – | – | – |
| GRU | **10** | 8 | 1 | 0 | **8900** | 23300 | 48600 | – | 7380 | 28800 | 67500 | – |
| S-LSTM | 1 | 1 | 0 | 0 | 22800 | 38100 | – | – | 13400 | 33500 | – | – |
| MGU | 5 | 4 | 0 | 0 | 11000 | 35800 | – | – | **6730** | 31600 | – | – |
| eGRU | 6 | 3 | 3 | 6 | 27500 | 34900 | 36500 | 36200 | 15700 | 28800 | 34600 | 38100 |
| SGU | 0 | 0 | 0 | 0 | – | – | – | – | – | – | – | – |
| IndRNN | 0 | 0 | 0 | 0 | – | – | – | – | – | – | – | – |
| YamRNN | 1 | **10** | **10** | **10** | 26300 | **7780** | **5070** | **4970** | 11900 | **4700** | **3410** | **3670** |

**Table 6.**
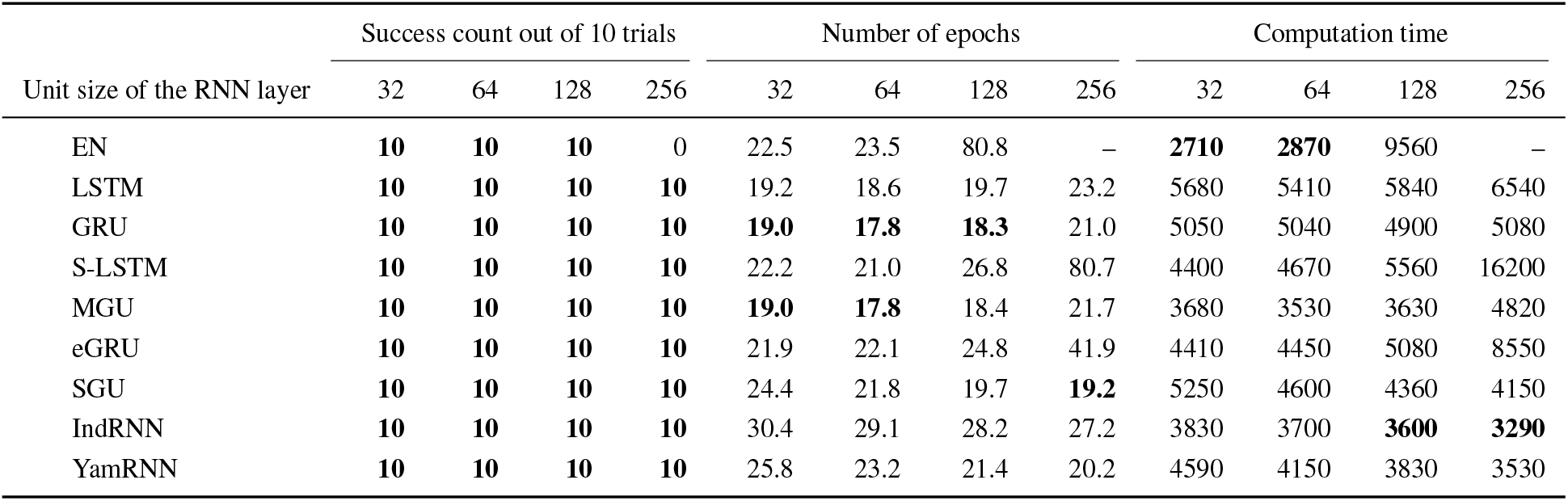
Results on test dataset 4 (IMDb), where the input sequence lengths differ, ranging from 11 to 2494. The number of epochs and computation time (seconds) are the mean values of the successful trials. The best values for each category are shown in bold. Three significant digits are displayed.

| Unit size of the RNN layer | Success count out of 10 trials |  |  |  | Number of epochs |  |  |  | Computation time |  |  |  |
| --- | --- | --- | --- | --- | --- | --- | --- | --- | --- | --- | --- | --- |
|  | 32 | 64 | 128 | 256 | 32 | 64 | 128 | 256 | 32 | 64 | 128 | 256 |
| EN | **10** | **10** | **10** | 0 | 22.5 | 23.5 | 80.8 | – | **2710** | **2870** | 9560 | – |
| LSTM | **10** | **10** | **10** | **10** | 19.2 | 18.6 | 19.7 | 23.2 | 5680 | 5410 | 5840 | 6540 |
| GRU | **10** | **10** | **10** | **10** | **19.0** | **17.8** | **18.3** | 21.0 | 5050 | 5040 | 4900 | 5080 |
| S-LSTM | **10** | **10** | **10** | **10** | 22.2 | 21.0 | 26.8 | 80.7 | 4400 | 4670 | 5560 | 16200 |
| MGU | **10** | **10** | **10** | **10** | **19.0** | **17.8** | 18.4 | 21.7 | 3680 | 3530 | 3630 | 4820 |
| eGRU | **10** | **10** | **10** | **10** | 21.9 | 22.1 | 24.8 | 41.9 | 4410 | 4450 | 5080 | 8550 |
| SGU | **10** | **10** | **10** | **10** | 24.4 | 21.8 | 19.7 | **19.2** | 5250 | 4600 | 4360 | 4150 |
| IndRNN | **10** | **10** | **10** | **10** | 30.4 | 29.1 | 28.2 | 27.2 | 3830 | 3700 | **3600** | **3290** |
| YamRNN | **10** | **10** | **10** | **10** | 25.8 | 23.2 | 21.4 | 20.2 | 4590 | 4150 | 3830 | 3530 |

**Table 7.** Results on test dataset 5 (Reuters), where the input sequence lengths differ, ranging from 13 to 2376. The number of epochs and computation time (seconds) are the mean values of the successful trials. The best values for each category are shown in bold. Three significant digits are displayed.

| Unit size of the RNN layer | Success count out of 10 trials |  |  |  | Number of epochs |  |  |  | Computation time |  |  |  |
| --- | --- | --- | --- | --- | --- | --- | --- | --- | --- | --- | --- | --- |
|  | 32 | 64 | 128 | 256 | 32 | 64 | 128 | 256 | 32 | 64 | 128 | 256 |
| EN | 4 | 9 | **10** | 0 | 375 | 126 | 101 | – | 6790 | 3190 | 3820 | – |
| LSTM | **10** | 9 | **10** | **10** | 246 | 235 | 194 | 207 | 20600 | 16900 | 16100 | 16200 |
| GRU | **10** | **10** | **10** | **10** | 156 | 81.2 | 50.5 | 49.2 | 11300 | 5670 | 3710 | 3410 |
| S-LSTM | 6 | **10** | **10** | 8 | 322 | 135 | 108 | 298 | 13000 | 8640 | 7040 | 15500 |
| MGU | **10** | **10** | **10** | **10** | 206 | 177 | 57.4 | 47.0 | 12700 | 11200 | 3540 | 2860 |
| eGRU | **10** | **10** | **10** | **10** | 238 | 159 | 111 | 160 | 14800 | 9930 | 6940 | 9820 |
| SGU | **10** | **10** | **10** | **10** | 96.4 | 60.2 | 46.8 | 39.1 | 6720 | 4210 | 3260 | 2630 |
| IndRNN | **10** | **10** | **10** | **10** | 144 | 115 | 91.6 | 74.7 | 5350 | 4320 | 3470 | 2760 |
| YamRNN | **10** | **10** | **10** | **10** | **45.0** | **35.3** | **29.4** | **24.8** | **2670** | **2090** | **1840** | **1520** |

Collectively, as shown in the five test experiments, the learning speed of YamRNN was superior to those of the existing methods if the number of parameters was set to be large enough. The learning speed of the existing RNNs decreased depending on the increase in the number of parameters, resulting in unsuccessful completion of computation on test datasets 2 and 3. Taking these properties of YamRNN and the other RNNs into consideration, for practical use, choosing YamRNN would be an option when existing RNNs with more parameters could not complete their learning process within practical time.

### 4.4 Convergence performance of YamRNN

In the previous section, we compared the performance of YamRNN to the existing RNNs from the perspective of learning speed. We here reiterate that the objective of the study was to develop a compact architecture with a higher learning speed than large architectures. In this regard, we succeeded in achieving the goal, as shown in the previous section. In this section, we additionally examined the performance of RNNs; we compared the convergence performance of YamRNN to that of the existing RNNs. To this end, we computed the maximum accuracy, which each RNN reached when the number of epochs in the learning process was limited to 1000, 3000, or 5000. For this experiment, we used SP500–100 because with this dataset the RNNs were expected to converge within practical time. Furthermore, this dataset is not as difficult as SP500–300, SP500–500, or Reuters, nor is it as easy as IMDb. Thus, we could expect to examine the differences in the performance of each RNN properly. As shown in Table 8, the accuracy of MGU was excellent when the parameter size was small (256) and the number of epochs was 1000 or 3000. This result is consistent with that of Table 3. Conversely, when the number of epochs was large (5000), the accuracy of GRU was the best. Because the parameter size of GRU was larger and thus of higher expressive power than the other existing RNNs except LSTM, GRU could achieve the highest accuracy when the number of epochs was sufficiently large. However, considering the fact that LSTM, which had larger parameters and thus could be expected to have more expressive power, could not achieve a higher accuracy, the structure of the core algorithm is more important than the size of the parameters to obtain higher convergence performance. In this experiment, when the parameter size was small, the performance of YamRNN was not good regardless of whether the number of epochs was 1000 or 5000. The low convergence performance would be one of the weak points of YamRNN. On the other hand, when the parameter size was large (1024), the performance of almost all the RNNs deteriorated compared to the smaller parameters and that of YamRNN was the best among all them; this result is consistent with the result in Table 3. In particular, the maximum accuracy of YamRNN (0.970), which YamRNN reached when the number of epochs was 5000, was far superior to all values achieved by the other RNNs. Regarding the computation time, IndRNN always had the best performance because the parameter size of IndRNN was simply the smallest among the RNNs. However, the convergence performance of IndRNN was not good in all cases; thus, this excellent learning speed is not useful practically. At the same time, the computation time of YamRNN was also excellent, especially for large parameter sizes. Although the convergence performance of YamRNN with smaller parameter sizes was not good, as previously mentioned, this excellent computation speed would compensate for the weak point of YamRNN. For example, whereas GRU with smaller parameter sizes took 1670 seconds to reach the accuracy of 0.882, which was the best score achieved by GRU in all experiments, YamRNN took only 1474 seconds to reach an accuracy of 0.970 with larger parameter values. More specifically, because YamRNN has the property that its architecture is compact and its convergence performance does not deteriorate with increasing parameter size, YamRNN can display good learning performance (speed and convergence) by increasing its parameter size.

**Table 8.** Comparison of convergence performance on test dataset 1 (SP500–100). The accuracy and computation time (seconds) is the mean value of the 10 trials. The best values for each category are shown in bold. Three significant digits are displayed.

| Number of epochs | Small (256) |  |  |  |  |  | Large (1024) |  |  |  |  |  |
| --- | --- | --- | --- | --- | --- | --- | --- | --- | --- | --- | --- | --- |
|  | Accuracy |  |  | Computation time |  |  | Accuracy |  |  | Computation time |  |  |
|  | 1000 | 3000 | 5000 | 1000 | 3000 | 5000 | 1000 | 3000 | 5000 | 1000 | 3000 | 5000 |
| EN | 0.545 | 0.602 | 0.687 | 120 | 357 | 593 | 0.531 | 0.533 | 0.534 | 206 | 614 | 1021 |
| LSTM | 0.572 | 0.710 | 0.784 | 411 | 1228 | 2046 | 0.561 | 0.662 | 0.829 | 760 | 2278 | 3797 |
| GRU | 0.589 | 0.772 | **0.882** | 335 | 1002 | 1670 | 0.547 | 0.586 | 0.812 | 610 | 1827 | 3044 |
| S-LSTM | 0.576 | 0.681 | 0.752 | 247 | 739 | 1230 | 0.548 | 0.596 | 0.784 | 440 | 1317 | 2195 |
| MGU | **0.608** | **0.773** | 0.861 | 253 | 759 | 1265 | 0.534 | 0.536 | 0.558 | 449 | 1343 | 2238 |
| eGRU | 0.562 | 0.648 | 0.704 | 236 | 706 | 1176 | 0.578 | 0.762 | 0.900 | 429 | 1281 | 2134 |
| SGU | 0.535 | 0.538 | 0.544 | 276 | 824 | 1372 | 0.542 | 0.558 | 0.585 | 475 | 1419 | 2364 |
| IndRNN | 0.533 | 0.533 | 0.533 | **106** | **315** | **524** | 0.533 | 0.533 | 0.534 | **123** | **363** | **604** |
| YamRNN | 0.552 | 0.643 | 0.728 | 185 | 553 | 924 | **0.608** | **0.887** | **0.970** | 296 | 885 | 1474 |

### 4.5 Investigating the topology of YamRNN

As shown in Tables 3 to 7, the learning speed of YamRNN continued to increase depending on the increase of parameter size. Whereas the learning speed of the existing RNNs increased or decreased with increasing parameter size depending on the situation, only YamRNN consistently increased the learning speed with increasing parameter size. This phenomenon could reflect the fundamental differences between YamRNN and the other architectures, caused by the difference in topology of the core architectures. At least, the high performance of YamRNN was not just because of the smaller number of parameters. In the first three test problems, the number of parameters of GRU on 512 units was 16896 and that of YamRNN on 1024 units was 17408. As shown in Table 4, the learning speed of GRU on 512 units (9520 seconds) was clearly different from that of YamRNN on 1024 units (1770 seconds), even though the number of parameters was almost the same in both the architectures. While S-LSTM is a derivative of LSTM and MGU, eGRU and SGU are derivatives of GRU, YamRNN was developed in a completely different way. The biggest difference between YamRNN and the existing architectures lies in the topology implied by the last equation of each architecture for computing output and memory vectors, **h**_*t*_. It is one of the important results of the study that we can clarify that topology contributes to the difference in performance. In order to study it more, we attempted to elucidate the properties of the topology of YamRNN. To this end, we compared the performance (learning speed) of derivatives of YamRNN and MGU on test dataset 1 (SP500–100). We chose MGU for comparison because its architecture is similar to that of YamRNN, although their learning performance differs; for example, the first equation of MGU (equation 2.16) and that of YamRNN (equation 4.1) are the same, the second equation of MGU (equation 2.17) and that of YamRNN (equation 4.2) use a hyperbolic tangent function. Further, the learning performance of MGU differs completely from that of YamRNN, as shown in Table 3, where the learning speed of YamRNN improves with increasing parameter size and the behavior is opposite for MGU. To generate derivatives of the RNNs, we swapped terms of the RNNs. For simplicity of presentation, we define the following equations:

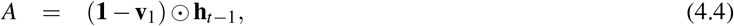

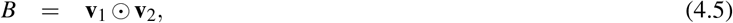

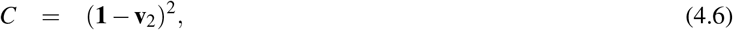

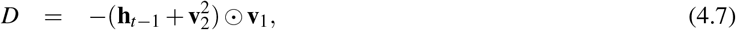

where A and B are derived from the third equation of MGU (equation 2.18), and C and D are derived from the third equation of YamRNN (equation 4.3). As the notable difference between MGU and YamRNN lies in their third equations, which both calculate **h**_*t*_, we removed the terms from the third equation and added these terms to the third equation. Here, we use the notation “MGU – A + C” to mean the term A is removed from and the term C is added to the third equation of MGU. The number of possible derivatives is eight: “MGU – A + C”, “MGU – A + D”, “MGU – B + C”, “MGU – B + D”, “YamRNN – C + A”, “YamRNN – C + B”, “YamRNN – D + A” and “YamRNN – D + B”. For these derivatives of RNNs, we conducted benchmarking on SP500–100 in the same manner as the experiment in Table 3. At this time, we identified the maximum accuracy each unsuccessful RNN achieved during 50000 epochs. We examined this information to help us understanding the performance of each RNN, because most of the RNNs could not complete the computation within the threshold number of epochs. As shown in Table 9, among the derivatives, only “YamRNN – C + B” completed the trials. Although the learning speed of the derivative was not better than the original RNNs, we confirmed that the learning speed improved with increasing parameter size. In addition, according to the maximum accuracy, the performance of “YamRNN – C + A” and “MGU – B + D” appeared to improve with increasing parameter size. The common property of these architectures is that they had the term D. Thus, the term D might be important for better performance. However, “MGU – A + D” also had the term D but its performance was poor; therefore, this theory does not always hold and is not beyond speculation. The clear fact obtained by the experiment is that the behavior of “YamRNN – C + B” was similar to YamRNN. The third equation of “YamRNN – C + B” is defined as follows:

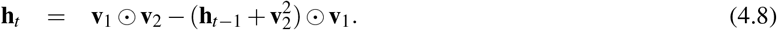

**Table 9.**
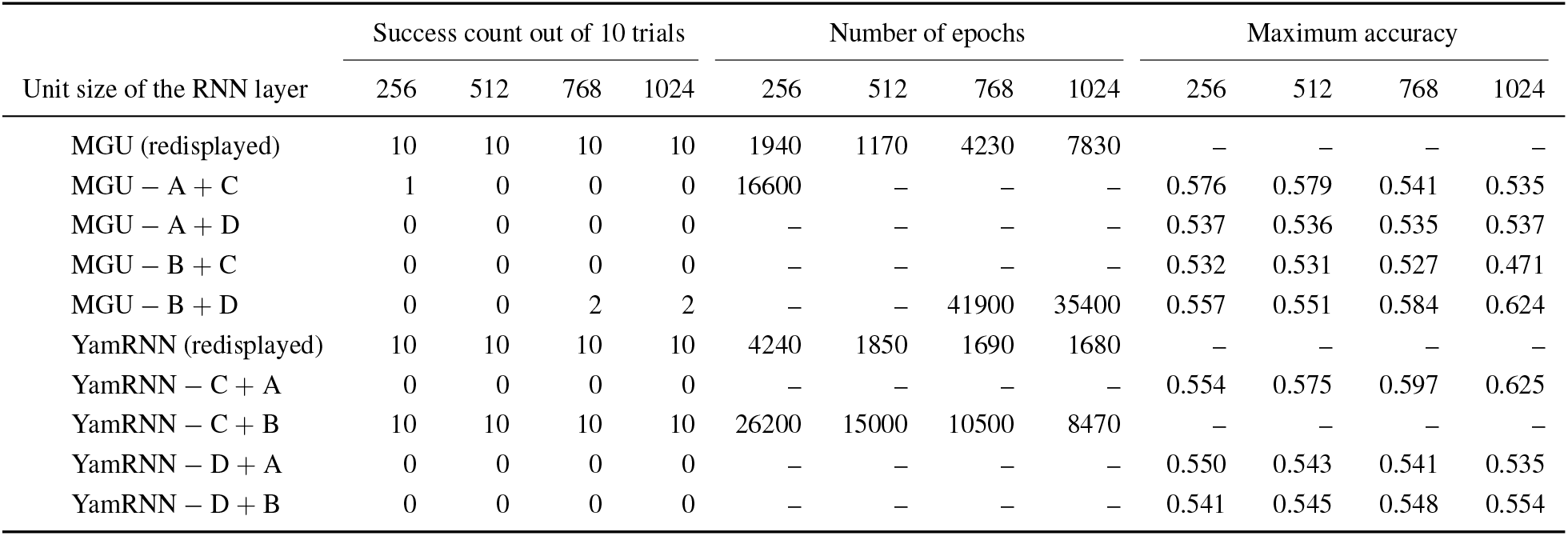
Comparison of derivatives of YamRNN and MGU on test dataset 1 (SP500–100). The number of epochs is the mean value of the successful trials and maximum accuracy is the mean value of the unsuccessful trials. Three significant digits are displayed. Note that “−X” or “+ X” denotes the term X is removed from or added to the third equation (equations 2.18 or 4.3) of the RNNs. The results of the original MGU and YamRNN are redisplayed from Table 3.

As shown, “YamRNN – C + B” and YamRNN differ from the other RNNs in terms of their having the squared term of the intermediate vector, 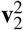. Next, in order to examine the effect of the squared term on the performance, we generated two derivatives. The first derivative was generated from MGU by replacing **v**_2_ in equation 2.18 with 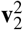, and the second was generated from YamRNN by replacing 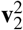 in equation 4.3 with **v_2_**. We denoted these RNNs as 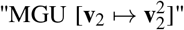 and 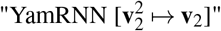, respectively. The third equation of 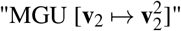 is defined as follows:

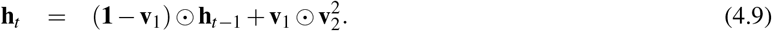

In contrast, the third equation of 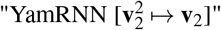 is defined as follows:

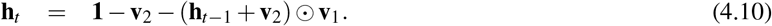

Note that the first term of equation 4.3 was expanded and subsequently replaced with **v_2_**, resulting in **1** – **v_2_** rather than (**1** – **v_2_**)^2^. For these derivatives of RNNs, we conducted benchmarking on SP500–100 in the same manner as the experiment in Table 3. As shown by the results in Table 10, the performance of neither RNN improved compared to the original RNNs. Further, the behavior of 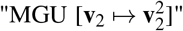 was not similar to YamRNN, even though the derivative included the squared term. On the other hand, the performance of 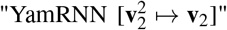 continued to improve with increasing parameter size, even though the derivative did not include the squared term. These facts suggest that the squared term is not the key factor to realize the unique property of YamRNN. Collectively, from the results of this series of experiments, we reconfirmed that the performance of RNNs is stipulated not only by elements in the RNNs but also by the complex combination of those elements. In addition to the means of generating the existing compact RNNs, we discovered YamRNN from the vast RNN space without any premises and YamRNN has a unique property with a unique topology. From this perspective, our strategy to develop RNNs would be valid for developing RNNs with novel properties. In the study, we could not completely clarify the property of the topology of YamRNN and we cannot discuss it further because there are only a few clues for further consideration. This issue will be investigated in future study.

**Table 10.** Comparison of derivatives of YamRNN and MGU on test dataset 1 (SP500–100). The accuracy and computation time (seconds) are the mean values of the 10 trials. Three significant digits are displayed. The results of original MGU and YamRNN are redisplayed from Table 3.

| Unit size of the RNN layer | Success count out of 10 trials |  |  |  | Number of epochs |  |  |  | Computation time |  |  |  |
| --- | --- | --- | --- | --- | --- | --- | --- | --- | --- | --- | --- | --- |
|  | 256 | 512 | 768 | 1024 | 256 | 512 | 768 | 1024 | 256 | 512 | 768 | 1024 |
| MGU (redisplayed) | 10 | 10 | 10 | 10 | 1940 | 1170 | 4230 | 7830 | 491 | 421 | 1750 | 3480 |
| MGU [$\mathbf{v}_2 \mapsto \mathbf{v}_2^2$] | 10 | 10 | 10 | 10 | 8940 | 4150 | 7500 | 7240 | 2310 | 1520 | 3150 | 3280 |
| YamRNN (redisplayed) | 10 | 10 | 10 | 10 | 4240 | 1850 | 1690 | 1680 | 795 | 461 | 473 | 507 |
| YamRNN [$\mathbf{v}_2^2 \mapsto \mathbf{v}_2$] | 10 | 10 | 10 | 10 | 12800 | 5000 | 3880 | 3610 | 2300 | 1170 | 1050 | 1050 |

### 4.6 Summary of the performance of YamRNN and future study

One of the major advantages of YamRNN could be its scalability against parameter increase. At least, YamRNN could reach the answer with the large parameter settings for which LSTM and GRU failed during computation. From this perspective, YamRNN was useful and could be an option for learning contextual information. However, we must note that we do not claim YamRNN is the best possible architecture, only that it will be a useful option for problems including contextual information. The most essential point related to machine learning applications is that there is no architecture which always the best performance for arbitrary problems. This was illustrated by our test results, where the best architecture changed depending on parameter settings such as data length and the number of units in the layer. When doing machine learning analysis, we must choose the optimal architecture suitable for problems by trial and error. Although in the study we could not analyze for what type of problem YamRNN was most suited, it was clear that YamRNN was compact and thus had an advantage in learning speed. Further advantages and weak points of YamRNN could be revealed by applying YamRNN to various real world problems. Finally, this study has the limitation that we could not investigate all possible architectures by our stochastic search method. In the study, we generated 30000 RNNs by combining operators and operands randomly. The RNNs were restricted to having only two intermediate equations (**v**_1_ and **v**_2_). This means in principle that there existed the possibility that the existing RNN with two intermediate equations such as MGU could be regenerated as a candidate RNN. However, MGU and other existing RNNs were not generated in the 30000 RNNs. This fact suggests that the extent of generating RNNs in the study is not sufficient. As we randomly generated RNNs from the vast RNN space, we believe the size of the search space is sufficient. On the other hand, the density of the generated RNNs in the search space, namely, the sampling number, may have been insufficient. If we expand the search range, there may be architectures with higher performance than YamRNN. However, at the same time, this limitation could be considered as a chance for further improvement. To find better architectures, search optimization methods such as Bayesian optimization and evolutionary algorithm could be useful.

## 5 Conclusion

In this study, we developed a novel RNN architecture, YamRNN, with a compact structure. We examined the performance of the RNN and confirmed that it provides normal learning performance for sequence data and excellent learning speed. The number of parameters of YamRNN was about 50% and 67% that of LSTM and GRU, respectively. The learning times required to achieve the same learning performance as LSTM and GRU were shortened at maximum by 85% and 80%, respectively, on a sequence classification task. One of the most outstanding advantages of YamRNN is its scalability against an increase in the number of parameters. This novel RNN architecture is expected to be useful for addressing various problems involving sequential data, such as speech recognition and generation, language translation, and image and text analysis. In addition, our success in developing this novel architecture would suggest that there is room for designing even better architectures, because we did not examine all feasible architectures in this study.

## Acknowledgement

All computations were performed on the NIG supercomputer at ROIS National Institute of Genetics.

## Availability

The code of the RNNs for Theano and TensorFlow, and the test datasets constructed in the study are available on GitHub, https://github.com/yamada-kd/YamRNN.git.

## Notes

This work was supported in part by the Top Global University Project from the Ministry of Education, Culture, Sports, Science, and Technology of Japan (MEXT), KAKENHI from the Japan Society for the Promotion of Science (JSPS) under Grant Number 18K18143 and 19J00950, and Platform Project for Supporting Drug Discovery and Life Science Research (Basis for Supporting Innovative Drug Discovery and Life Science Research (BINDS)) from AMED under Grant Number JP20am0101067.

### Competing Interest Statement

The authors have declared no competing interest.

### Summary of Updates

Tables 6-10 updated.

